# Elasticity of Whole Blood Clots Measured via Volume Controlled Cavity Expansion

**DOI:** 10.1101/2023.02.17.528966

**Authors:** Hannah Varner, Gabriella P. Sugerman, Manuel K. Rausch, Tal Cohen

**Affiliations:** Department of Mechanical Engineering, Massachusetts Institute of Technology, Cambridge, 02139, MA, USA; Department of Biomedical Engineering, University of Texas at Austin, Austin, 78712, TX, USA; Department of Aerospace Engineering and Engineering Mechanics, University of Texas at Austin, Austin, 78712, TX, USA; Oden Institute for Computational Engineering and Sciences, University of Texas at Austin, Austin, 78712, TX, USA; Department of Civil and Environmental Engineering, Massachusetts Institute of Technology, Cambridge, 02139, MA, USA

**Author notes:** Corresponding author. MIT Civil and Environmental Engineering. 77 Massachusetts Avenue Cambridge, MA, 02139.

**Keywords:** Blood clot, Cavity expansion, Ogden model, Hyperelasticity

## Abstract

Measuring and understanding the mechanical properties of blood clots can provide insights into disease progression and the effectiveness of potential treatments. However, several limitations hinder the use of standard mechanical testing methods to measure the response of soft biological tissues, like blood clots. These tissues can be difficult to mount, and are inhomogeneous, irregular in shape, scarce, and valuable. To remedy this, we employ in this work Volume Controlled Cavity Expansion (VCCE), a technique that was recently developed, to measure local mechanical properties of soft materials in their natural environment. Through a highly controlled volume expansion of a water bubble at the tip of an injection needle, paired with simultaneous measurement of the resisting pressure, we obtain a local signature of whole blood clot mechanical response. Comparing this data with predictive theoretical models, we find that a 1-term Ogden model is sufficient to capture the nonlinear elastic response observed in our experiments and produces shear modulus values that are comparable to values reported in the literature. Moreover, we find that bovine whole blood stored at 4°C for greater than 2 days exhibits a statistically significant shift in the shear modulus from 2.53 ± 0.44 kPa on day 2 (*N* = 13) to 1.23 ± 0.18 kPa on day 3 (*N* = 14). In contrast to previously reported results, our samples did not exhibit viscoelastic rate sensitivity within strain rates ranging from 0.22 – 21.1 s^−1^. By surveying existing data on whole blood clots for comparison, we show that this technique provides highly repeatable and reliable results, hence we propose the more widespread adoption of VCCE as a path forward to building a better understanding of the mechanics of soft biological materials.

**Graphical Abstract:** 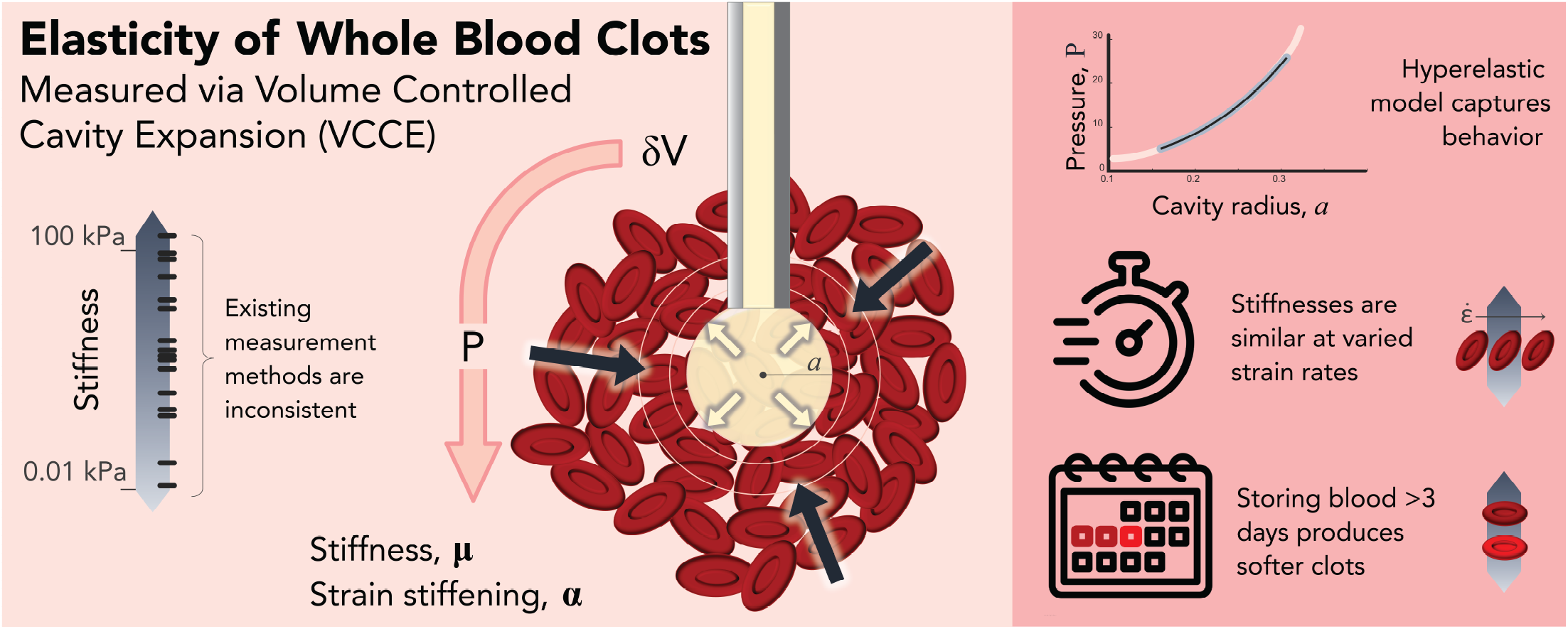

**Highlights:**

- Volume controlled cavity expansion overcomes common obstacles to testing biological samples
- Whole blood clot elasticity is well captured by the Ogden hyperelastic material model
- Shear modulus strain-rate sensitivity was not observed in clots for moderate rates

## 1. Introduction

While blood clot coagulation is vital for human survival, thrombi are also a leading cause of mortality and morbidity world wide (World Health Organization, 2020). Formed naturally following a traumatic injury, after surgery, or even during pregnancy, thrombus dislodgement can be fatal, leading to heart attack, stroke, and pulmonary embolism. The mechanical stiffness and relaxation properties of whole blood clots, and of contracted thrombi in vivo, have been highlighted as potentially indicative of disease progression, and are thus crucial for the development of methods for prevention and treatment (Litvinov and Weisel, 2016; Hinnen et al., 2005; Liebeskind et al., 2011; Marder et al., 2006).

Blood clots are complex, hierarchical biological polymers composed of long chains of fibrin and interspersed with platelets and whole blood cells. The prevalence of cells within the structure means that characterization of the mechanical stiffness of isolated constituents is insufficient to capture the response of a whole clot as it might exist in the body as mechanical loading changes the cell shapes and packing (Tutwiler et al., 2018; Liang et al., 2017; Gersh et al., 2009). In contrast to fibrin or platelet poor clots, whole blood clots containing red blood cells are less stiff and demonstrate different viscoelastic behavior (Gersh et al., 2009; Huang et al., 2013; Riha et al., 1999; Litvinov and Weisel, 2017; Tynngård et al., 2006).

The conditions required to dislodge a clot are dependent on not only the geometric boundary conditions, but also the mechanical properties of the clot such as stiffness and viscoelastic relaxation. However, quantification of mechanical properties in soft biological tissues remains challenging. Conventional mechanical testing methods often necessitate homogeneous, macroscopic specimens of predefined geometry (e.g. dog-bone in tension testing, or plugs in compression testing), or may only access bulk properties near the surface of a sample (e.g. indentation testing), and are incapable of mapping spatial variation within a material (e.g. shear rheology). The heterogeneity of most tissues, combined with the finding that mechanical properties can change significantly when a tissue is removed from its native environment (Nickerson et al., 2008; Zimberlin et al., 2010), necessitate a mechanical testing technique that can be used to probe multiple locations of these heterogenous tissues *in vivo*. The variability and range of local mechanical properties can be used to quantify disease progression or the safe limits of mechanical loading that human tissues can experience.

The study of mechanical properties in thrombus mimics largely bares out the complications of biological tissue testing described above. Consensus has been growing in recent liteature that whole blood clots are strain-stiffening (Riha et al., 1999; Krasokha et al., 2010; Malone et al., 2018; Sugerman et al., 2020; Chueh et al., 2011). However, the quantification of stiffness within whole clots varies widely across literature. While some of the variation can be attributed to differences in the individual donor animals, a principal complication of obtaining material properties for these soft and highly hydrated tissue is the challenge in executing standard mechanical tests. To avoid the irregular shape of clots formed *in vivo*, researchers have turned to clots formed in a laboratory setting with more repeatable macro-scale geometry and homogeneity but have experienced mixed success in crafting and successfully testing these “standard” specimens (Malone et al., 2018; Krasokha et al., 2010). Authors have performed mechanical characterization of whole clots derived from human, porcine and bovine blood using tensile elongation (Malone et al., 2018; Krasokha et al., 2010), pure shear (Sugerman et al., 2020, 2021b), compression or indentation testing (Litvinov and Weisel, 2016; Liang et al., 2017; Malone et al., 2018; Chueh et al., 2011; Weafer et al., 2019; Johnson et al., 2021), and oscillatory shear rheometry (Gersh et al., 2009; He et al., 2022; Malone et al., 2018; Riha et al., 1999; Tynngård et al., 2006; van Kempen et al., 2016).

However, these studies report elasticity values spanning three orders of magnitude. These variations may be attributed to some methods probing the bulk, while others test the local properties, the difficulty in applying standard geometries to samples, and the strain stiffening nature of biological materials. Furthermore, values are frequently reported from a small number of replicates given the difficulties in executing the tests and destructive nature of testing.

The idea of a needle-based method to measure mechanical properties of soft biological tissue has grown in popularity as a way to overcome the geometry, handling, and sample condition considerations. First Needle Induced Cavitation Rheology (NICR) (Zimberlin et al., 2007; Kundu and Crosby, 2009) and later Volume Controlled Cavity Expansion (VCCE) (Raayai-Ardakani et al., 2019a,b), allow for local material testing through recording the resistance of a sample to expansion of a cavity at the end of a needle. While NICR relies on the existence of a cavitation instability, highly strain-stiffening materials, such as blood clot, exhibit fracture and do not approach a cavitation limit. VCCE thus builds on a decade of NICR literature, while using an incompressible working fluid and volumetric control of the cavity, to access rich data from the full range of the material response (Raayai-Ardakani and Cohen, 2019; Nafo and Al-Mayah, 2021). Nafo and Al-Mayah (2021) demonstrate the flexibility of controlled cavity expansion testing to allow for selection of a constitutive model that is well matched to the tissue under test (i.e. porcine liver). Also using a strain-stiffening constitutive model, Mijailovic et al. (2021) show that VCCE captures material stiffness comparable to values from small strain microshear rheological testing in brain tissue, achieving this with high enough resolution to distinguish white matter from gray matter.

The precise volume control in VCCE allows for investigation of the rate-dependent properties of a sample. Both Mijailovic et al. (2021) and Ji et al. (2022) note that the stiffness values obtained during NICR tends to be correlate well with high strain rate testing. By employing VCCE at different strain rates, Chockalingam et al. (2021) were able to quantify the viscoelastic properties of Sylgard 184 polydimethylsiloxane (PDMS) samples.

In this work, we first propose the application of VCCE to testing of whole blood clots as a step towards *in vivo* tissue measurement. Second, we demonstrate the utility of VCCE in exploring the rate dependence of whole clot mechanical properties, which is of vital importance for enabling the application of computer simulation to the analysis of biological tissues such as clot in dynamic loading scenarios (Gasser et al., 2022). We find that while the elastic shear modulus does not exhibit sensitivity to loading strain rates in the range 0.22 - 21.1 s^−1^, the critical values (indicating an onset of damage) show appreciable rate sensitivity. Moreover, the storage time of the blood prior to coagulation significantly influences the measured mechanical response.

This paper is organized as follows: We begin by describing our experimental method and theoretical formulation in Section 2. Section 3 describes the results of our experiments, with presentations of the results organized by both the test rate (3.1) and storage time (3.2). We discuss our results in Section 4 and conclude in Section 5.

## 2. Materials and Methods

### 2.1. Whole blood clot

Whole blood clots were prepared from bovine blood collected with CPDA-1 anticoagulant at 14% volume anticoagulant/total volume and derived from a single donor animal (Lampire Biological Laboratories, PA,USA) (Sugerman et al., 2021a). The blood is stored at 4 °C prior to initiation of coagulation via the addition of calcium chloride to a final concentration of 20 mM. Between 20 and 25 mL of whole blood volume is placed in a 30 mL container using a serological pipette and the containers are covered to avoid water loss while they are incubated at 37 C for 90 minutes^1^. Testing is then performed within 90 minutes of removal from the incubator. After removing a blood clot sample from incubation, phosphate buffered saline (PBS) is added to the container such that the surface of the clot is fully covered to prevent drying and maintain osmotic balance.

### 2.2. VCCE experimental procedure

VCCE is used to initiate a metered expansion of the cavity. As in Chockalingam et al. (2021), the plunger of a 50*μL* syringe (Hamilton GASTIGHT of Reno, NV, USA) is controlled by an Instron Electropuls 3000 (Norwood, MA,USA) and the pressure is recorded by a PRESS-S-000 luer lock connected pressure sensor (PendoTech, Princeton, NJ, USA). We relate the displacement of the syringe plunger to the volumetric increase of the cavity as *V* = (4*/*3)*πa*^3^ where *a* is the effective radius of the cavity, assuming it is spherical. Raayai-Ardakani et al. (2019a) and Mijailovic et al. (2021) have independently shown that the error derived from non-spherical initial cavity is insignificant throughout the majority of the expansion.

A typical pressure-radius profile is shown schematically in Figure 1. The pressure increases during expansion of the cavity up until a critical radius *a*_*crit*_ at which point the sample fractures and pressure decreases dramatically from the peak at *P*_*crit*_ (Raayai-Ardakani et al., 2019b). Data within the dashed lines (*a*_*min*_ to *a*_*max*_) is used for fitting material parameters as discussed in Section 2.4.

**Figure 1:**
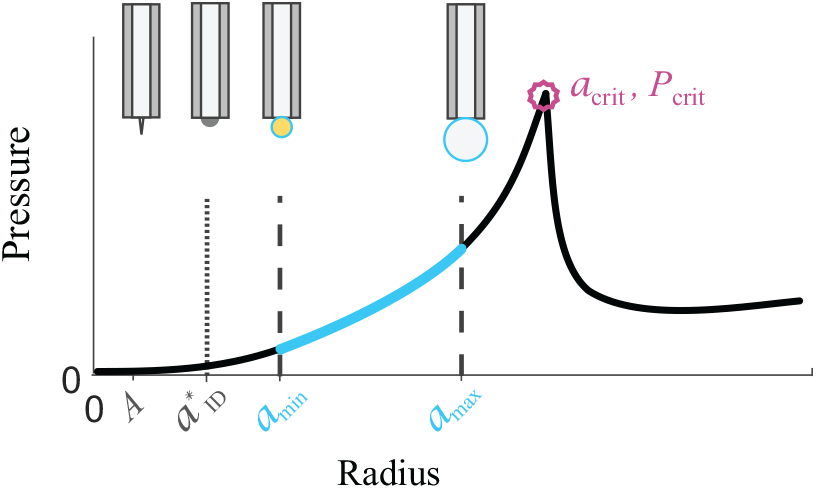
During a VCCE test the measured pressure increases as fluid is injected until a critical pressure *P*_*crit*_ is obtained, after which pressure decreases sharply as the sample fractures. A typical initial defect (modeled as a sphere with radius *A >* 0) is illustrated relative to the radius of a hemisphere defect the size of the needle inner diameter 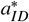. The bounds of elastic data used in fitting material properties are denoted by *a*_*min*_ and *a*_*max*_.

Prior to performing each experiment, we examine the water column to ensure that no air has entered the system, as this would lead to erroneous volume measurements. For pressure calibration we perform a mock injection into a container of the working fluid at the programmed test rate prior to material testing. The pressure from this step is subtracted from the experimental data to eliminate the contribution of viscous losses to the pressure.

The Instron is programmed to produce constant radial expansion rates 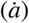 of 0.02, 0.08, 0.32, 0.64, and 1.60 mm/s, corresponding to elastic expansion times from 15 to 0.1875 s. The range is selected to include rates that have previously been used with VCCE to identify viscoelastic behavior of materials while the inertial effects of the cavity expansion can be neglected (Chockalingam et al., 2021; Cohen and Molinari, 2015).

When testing a clot, the needle is held stationary and the sample platform is raised until the needle first breaks the surface, then to a final depth approximately double the surface penetration depth, before “retracting” the sample slightly (see Figure 2A) in order to form an initial defect. Barney et al. (2019) have shown that retracting the needle during NICR reduces residual strain in the material, with a minimal effect on the measured mechanical properties. By strictly following this insertion protocol, we expect that the initial defects will be of a similar size in all the VCCE trials. Typical depths used for the whole blood clots presented here are shown in Figure 2A. All testing presented here uses a 25 gauge blunt needle (outer diameter ø_*OD*_=0.51 mm, inner diameter ø_*ID*_=0.26 mm).

**Figure 2:**
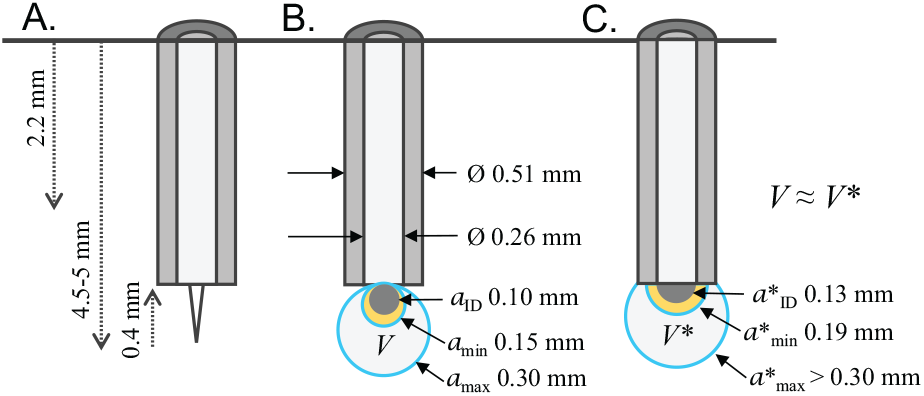
**A** provides dimensions of a typical insertion into the blood clot. Equivalent spherical radii used during data fitting are shown in **B**. These were informed by the geometry of the experimental set up described by **C**. For example, the yellow sphere in **B** with *a* = 0.153 mm would have the equivalent volume to a hemisphere with 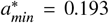 mm, the average of the inner and outer radii of the needle.

The working fluid for all tests is PBS with a small amount of water-based colorant (to enable visualization of test failures associated with fluid leaking from the cavity). Five minutes after insertion, or when the recorded pressure reaches 0 kPa, we initiate radial expansion from zero volume. One sample is tested at multiple locations, taking care to space each test away from previous tests or the container walls (*>*10X the diameter of the needle)

### 2.3. Theoretical foundation of VCCE

We employ the theoretical framework proposed in Raayai-Ardakani et al. (2019a) to relate the experimental results of the VCCE method to the material properties. We consider our test specimen to be an unbounded^2^ and incompressible body (an assumption justified for our experimental procedure in Section 4.4). A spherically symmetric internal cavity of undeformed radius *A* is embedded within the body and expanded to a new radius *a* by injecting an incompressible fluid that is assumed to be isobaric. With our assumption of spherical symmetry, the measured fluid pressure is directly related to the material response. We represent the material response using an arbitrary free energy function, which can be written in terms of the principle stretch components 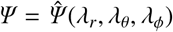, in the radial and circumferential directions, respectively. The circumferential stretch is *λ* = *λ*_*θ*_ = *λ*_*ϕ*_ = *r/R*, where *R* and *r* denote the radial coordinate in the undeformed and deformed configurations, respectively. Incompressibility implies *λ*_*r*_ = *λ*^−2^. Hence, the energy density can be simplified to write 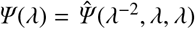.

To determine the pressure-stretch relationship, we first obtain the total elastic energy, *U*_*E*_, by integrating *Ψ* throughout the body. As shown by Raayai-Ardakani et al. (2019b), the cavity pressure can then be obtained directly by the differentiation *P* =−*∂U*_*E*_*/∂V*, where *V* represents the volume of the cavity. The resulting pressure can then be written in the form

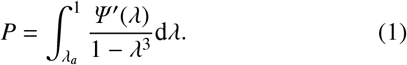

Pressure contributions from surface tension are not included in this formulation given that the working fluid creating the cavity and the material under test are both water-based.

In keeping with the observations of Sugerman et al. (2021b), Umale et al. (2013), Budday et al. (2017), and Lu et al. (2014) for thrombi mimics, spleen, brain and liver tissue, respectively, we use a 1-term Ogden model to capture the constitutive response of the whole blood clot and compare the measured values to existing literature. While this model is not without its limitations (Lohr et al., 2022), it is useful in the context of whole blood clots because it captures the full nonlinear strain stiffening response of the material. The 1-term Ogden model relates volumetric strain energy to a circumferential stretch *λ* in spherical coordinates as

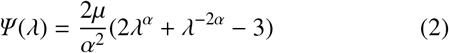

with the conventional shear modulus *μ* and strain stiffening parameter *α* (Ogden, 1972).

### 2.4. Material model fitting

Material parameters *μ* and *α*, and an effective initial cavity size *A* (used to calculate *λ*) are estimated using the native nonlinear least squares (NLLS) fitting algorithm within MATLAB. We first employ a multi-start method to fit a subset in order to determine appropriate ‘average’ initial conditions for the NLLS method. To ensure we have avoided a local minimum, we perform a sensitivity test to the initial conditions of the NLLS fit. Trials that did not converge on a global minimum with multiple seeds for initial radius (i.e. were highly sensitive to the initial guess) were excluded from the final data set; this occurred in *<* 10% of trials.

During the experiment, expansion begins from an initial defect of effective radius *A* and continues until the material fractures at *a*_*crit*_. Although the initial defect is not a perfect sphere, initial geometric imperfections have little effect on the pressure-volume response within the considered experimental range of data. This insensitivity was shown in two separate studies (Raayai-Ardakani et al., 2019a; Mijailovic et al., 2021) that, by employing computational simulations with various defect shapes, demonstrate that the deviation from the theoretical spherical solution is significant only for small expansion volumes. The spherical assumption is thus a good approximation for VCCE.

Accordingly, in the present experiments, the lower bound for fitting is chosen based on the system geometry: first calculating the volume of a hemisphere with *a*^*^ equal to the average of the needle outer and inner diameters ø_*OD*_ and ø_*ID*_, then finding the radius of the equivalent sphere containing this volume (*a*_*min*_ = 0.153 mm). The upper bound, *a*_*max*_ = 0.3 mm, is informed by the material response such that a majority of the trials have *a*_*crit*_ *> a*_*max*_. In the case that *a*_*crit*_ *< a*_*max*_, fitting was performed on the curve from *a*_*min*_ to *a*_*crit*_. A schematic of *a*_*min*_ and *a*_*max*_ is shown in Figure 2B and 2C. Note that fitting until *a*_*crit*_ in all trials results in an inverse dependence of *A* on the absolute value of *a*_*crit*_. This is contrary to the expectation that the size of the initial defect is independent of the loading conditions that follow after it is formed.

## 3. Results

To better examine trends in our results, we segment our data by expansion rate and by day (i.e. days of whole blood storage prior to clotting) when examining both the experimentally collected values and fitted material parameters. A selection of experimental data and the corresponding curve fits demonstrating the versatility of the Ogden model for our data can be seen in Section ESI 1 of the Electronic Supplemental Information.

### 3.1. Expansion rate

Table 1 and Figure 3 summarize our experimental results analyzed with respect to the radial expansion rate. Mean fitted values of the materials properties 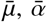 as well as the effective initial radius of the cavity, *Ā*, are provided (and denoted by the superimposed bar) along with the standard deviations (SD), for each specified expansion rate. For trials with 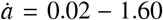 mm/s, stiffening is observed to be consistent across all rates considered (Figure 3E) with *α* = 5.76 0.27 (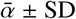 across all trials *N* = 22). Figure 3C shows that the initial defect radius *A* = 0.082 ± 0.013 mm is below 0.10 mm in all but one trial, indicating that the spherical volume of the initial defect is less than that of a hemispherical cap with the internal diameter of the needle^3^. We take this as an indication that the cavities created in our experiment are not simply expanding off of the face of the blunt needle (Figure 2C), but are in fact generating deformation that is resisted by the bulk of the material and can be well represented by a spherical field assumption (Figure 2B).

**Table 1:**
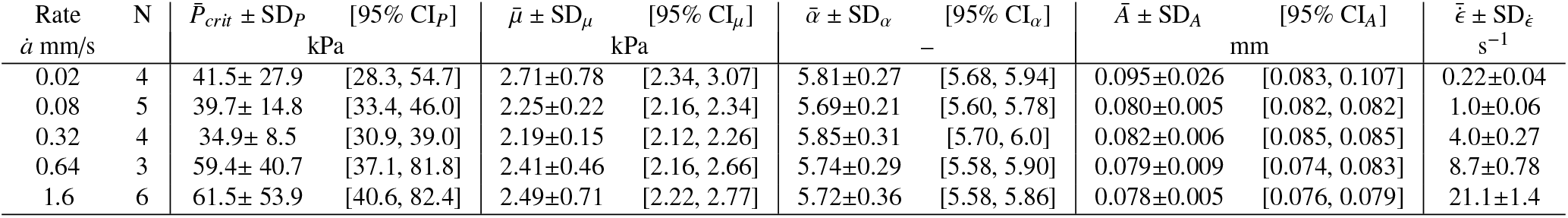
Results grouped by test rate. Clots from blood stored for ≤ 2 days.

**Figure 3:**
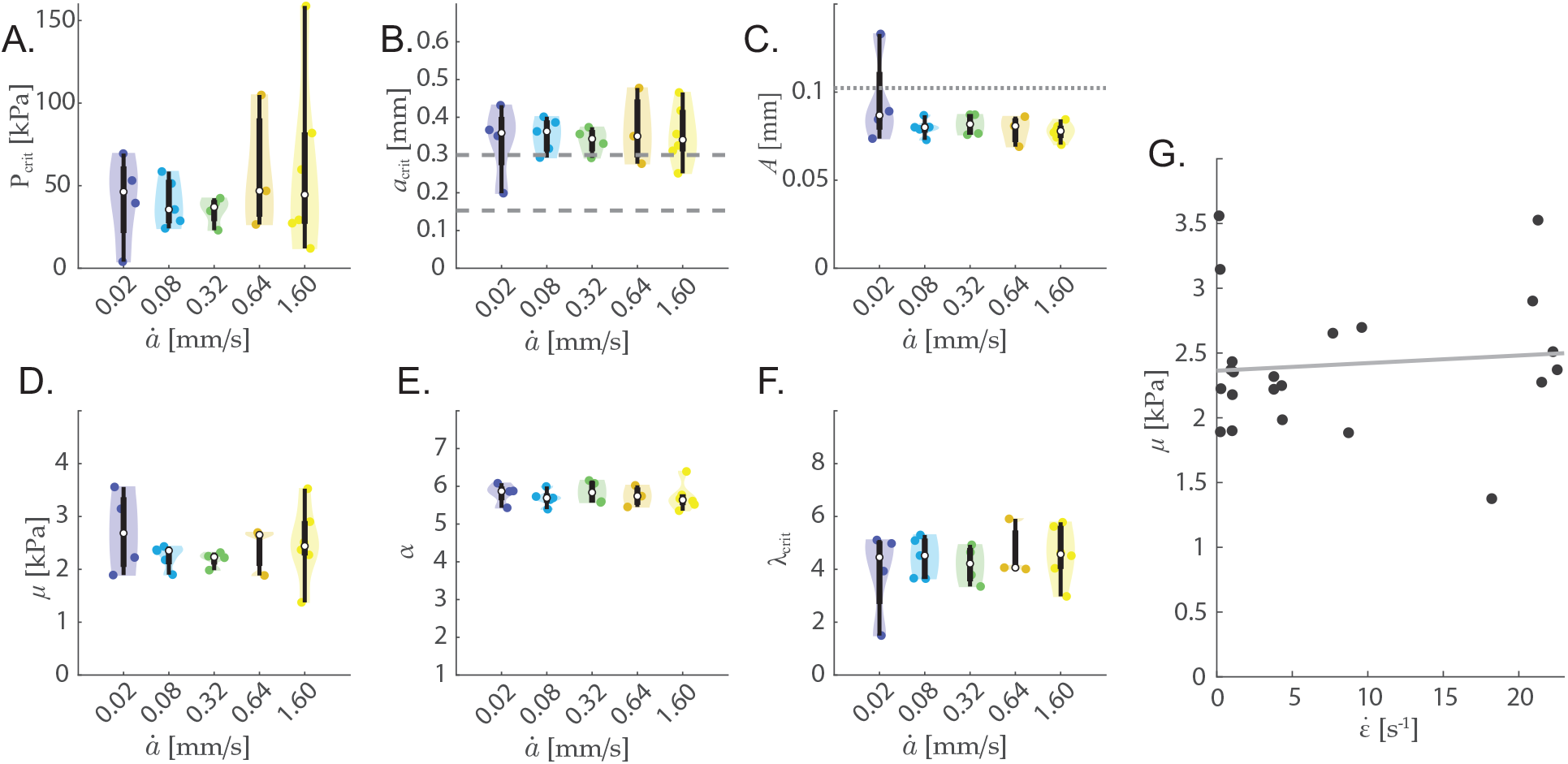
Experimental data grouped by rate for clots made from blood stored for ≤ 2 days. For **A**.-**F**. the median is marked on the plots by an open circle and the box extends from the first to third quartile. The dashed lines on **B**. indicate the region of data that was used during the NLLS fitting procedure. The dotted line on **C**. indicates the equivalent spherical radius for a hemispherical defect with the internal diameter of the needle. The fit material parameters (**D**. and **E**.) show no statistically significant differentiation. However, the measured modulus increases with increasing strain rate as shown in **G**. where a linear fit results in 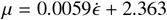 kPa with RMSD=0.53 kPa.

The experimental strain rate 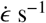 is determined from the ratio between the radial expansion rate from *a*_*min*_ to *a*_*max*_, and the fitted value of *A* for a given trial. Strain and stretch at the cavity wall relate as *ϵ* = *λ* − 1 resulting in 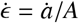. Accounting for the variability observed in each experiment results in strain rates from 0.22 ± 0.04 s^−1^ for 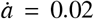 mm/s to 21.1 ± 1.4 s^−1^ for 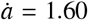 mm/s. Figure 3G relates *μ* to 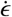 for each trial. Though not well described by a linear fit, with a root-mean-square deviation (RMSD) of 0.533 kPa for 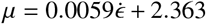 kPa, some variation with rate can be observed.

Employing an analysis of variance test (ANOVA) with *p <* 0.05 did not yield any statistically significant differentiation of material properties or critical values for the rates tested. The number of tests at each rate ranged from *N* = 3 to 6. *λ*_*crit*_ ranged from 1.50 to 5.91, with a mean of 4.36.

### 3.2. Blood storage time

Comparing all the trials performed in our experiments, we find that blood stored for more than 2 days produces a softer clot. An ANOVA test indicates a statistically significant decrease in the equilibrium shear modulus between each of the first two days *μ* = 2.24 kPa with a 0.95 confidence interval of [2.05, 2.43] kPa and 2.53 [2.41, 2.65] kPa, respectively, and the third day of testing 1.23 [1.19, 1.28] kPa. This is apparent in the raw pressure-radius curves shown in Figure 4G, as well as the stiffness values in 4D and Table 2.

**Figure 4:**
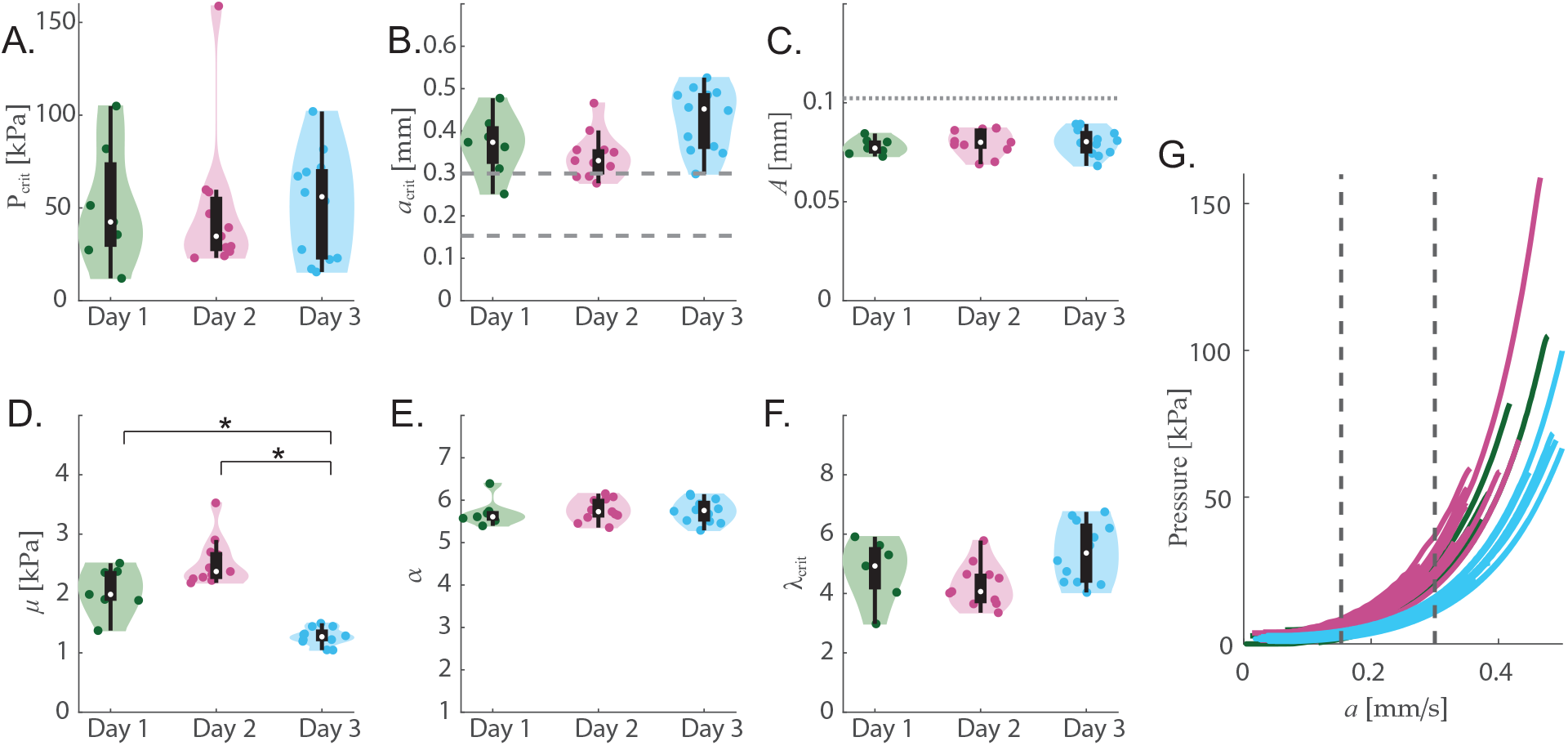
Experimental results grouped by days of storage show statistically significant variation in results for only the shear modulus *μ* (**D**.) Results marked with [ ]^*^ indicated statistical significance at p*<* 0.05. Median is marked by an open circle on all plots **A.-F**. and the box extends from the first to third quartile. The dashed lines on **B**. and **G**. indicate the region of data that was used during the NLLS fitting procedure. The dotted line on **C**. indicating the equivalent spherical radius for a hemispherical defect with the internal diameter of the needle. **G**. shows the full traces of experimental raw data collected. Traces are colored by test day with green indicating test performed *<*24 hours after collection, pink within 24 – 48 hours, and blue within 48 – 72 hours. Data after *P*_*crit*_ has been removed from the plot.

**Table 2:** Results grouped by days of storage show significant shift in shear modulus between 2 and 3 days. Mean and standard deviation include tests performed with 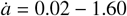 mm/s.

| Day | N | $\bar{\mu} \pm SD_{\mu}$<br>kPa | [95% CI$_{\mu}$] | $\bar{\alpha} \pm SD_{\alpha}$<br>– | [95% CI$_{\alpha}$] | $\bar{A} \pm SD_A$<br>mm | [95% CI$_A$] |
| --- | --- | --- | --- | --- | --- | --- | --- |
| 1 | 9 | $2.24 \pm 0.60$ | [2.05, 2.43] | $5.69 \pm 0.30$ | [5.6, 5.79] | $0.084 \pm 0.019$ | [0.078, 0.090] |
| 2 | 13 | $2.53 \pm 0.45$ | [2.41, 2.65] | $5.8 \pm 0.26$ | [5.73, 5.87] | $0.081 \pm 0.006$ | [0.079, 0.083] |
| 3 | 14 | $1.23 \pm 0.18$ | [1.19, 1.28] | $5.74 \pm 0.25$ | [5.68, 5.80] | $0.079 \pm 0.007$ | [0.078, 0.081] |

We see no statistically significant separation by days of blood storage for the experimental critical values *P*_*crit*_ (Figure 4A) and *a*_*crit*_ (4B), or *A* (4C), *α* (4E), and *λ*_*crit*_ (4F). Each test day includes data from multiple different blood clots created on that day from the same reservoir of donor blood but tested at variety of rates 0.02 mm/s - 1.60 mm/s. Including the 14 additional tests for blood stored for 3 days, the average and SD of *α* and *A* both remain consistent with SD *<* 15% of the value (*α* = 5.75 ± 0.26 and *A* = 0.081 ± 0.011 mm, *N* = 36).

## 4. Discussion

### 4.1. Elasticity values in context

It is instructive to compare the results obtained in this work with the reported results in the literature. The most commonly reported material parameter is the shear modulus. In Figure 5 we summarize reported shear moduli from 11 studies of whole blood clots using bovine, human, porcine, and ovine blood. The two most commonly used *in vitro* mechanical test methods, compression testing and rheology, result in some of the highest and lowest stiffness values for blood clot, respectively. Reported shear moduli in literature range from 0.013 to 16.8 kPa. With VCCE, we found a more narrow range of stiffness with *μ* = 1.38 to 3.53 kPa from *N* = 22 with an average of 2.41 kPa and SD of 0.52 kPa.

**Figure 5:**
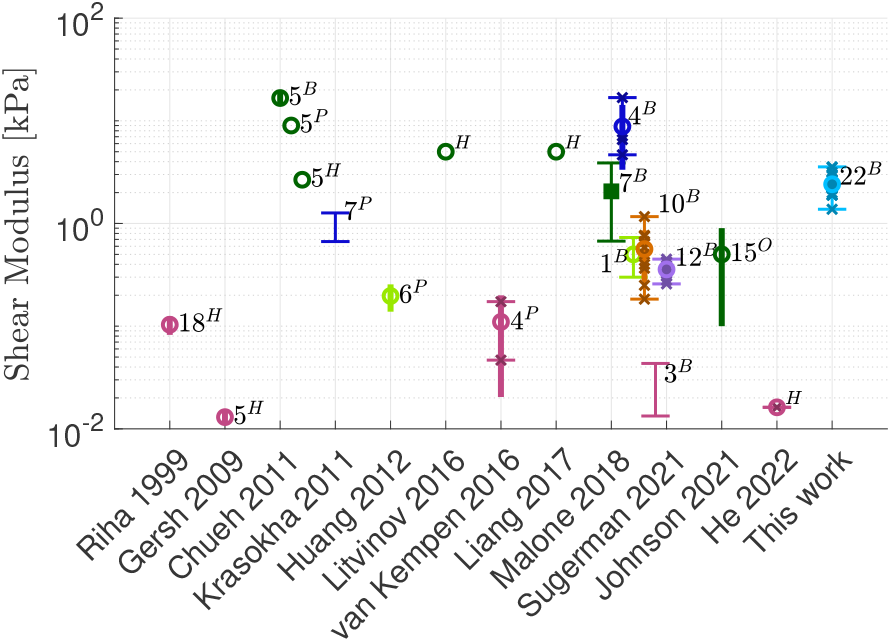
Comparison of literature results for whole blood clot elasticity testing. Tests are color coded by type of experiment performed. Dark green is compression, pink rheology, dark blue is tension, purple is simple shear, light green is elastography, orange is indentation and light blue is VCCE. If reported: ranges are indicated by thin lines with a horizontal bar at the end, the standard deviation is shown by a thick vertical line. Individual data points are crosses and open circles show the mean. Median is a filled square. The *N* value is written next to each data set and is the minimum value if authors reported *N* as a range. Blood for these test is from mixed origin (human, porcine, ovine, and bovine) each denoted by a superscript on the *N* value. Only lab-made, whole blood clots are considered.

Compared to the existing literature, VCCE enables more trials to be conducted on the same sample and provides repeatable results. Only one of the other studies in Figure 5 presents results with *N >* 15 and 75% present a modulus based of *N <* 10. In addition to the higher replicate count, VCCE is the only method of blood clot testing that allows for potential *in vivo* testing of clot samples since it does not require destruction of bulk tissue to extract a test specimen.

### 4.2. Rate insensitivity within test range

The consistency of *A* across rates and days validates the VCCE methodology. It is notable that despite a marked shift in stiffness after 2 days of storage, *Ā* is within 5 *μ*m across the three days. We take this as indication that the needle insertion process creates a consistent initial defect within the sample.

At the same time, the lack of distinct shear moduli for the rates tested is unexpected based on previous work in whole blood clots. Sugerman et al. (2020) documents a marginal increase in stiffness for pure shear tests conducted at strain rates of 2%s^−1^ vs. 50%s^−1^. In contrast, our experiments employed stretch rates between 22 and 2000%s^−1^ and show now increase in stiffness, possibly indicating a saturation in rate-dependent stiffening for whole blood clots. Further exploration of the rate dependence of *μ* is warranted to build a more complete picture of the slight increase in stiffness with strain rate (6 Pa per s^−1^) shown in Figure 3G and improve the prediction intervals of this trend.

As an alternative to our approach of examining the influence of expansion rate on the elastic modulus, VCCE could be used to perform relaxation tests as in Chockalingam et al. (2021) to obtain viscoelastic properties. Viscoelastic relaxation of blood clots has been considered by previous authors, but was not examined in this work. Sugerman et al. (2020) and Johnson et al. (2021) both use an exponential decay (Prony series) to model strain-dependent stress relaxation of whole clots. Meanwhile, van Kempen et al. (2016) propose a two term Maxwell model with 7 material parameters to capture the nonlinear viscous dissipation they observed in whole blood clots. By contrast, when studying aortic aneurysms, van Dam et al. (2008) propose a Leonov-type 3-term viscoelastic model that allows for nonlinear stress evolution that is fully characterized by two linear parameters of a spring-dash-pot type Maxwell model. A modified protocol from that presented here for whole blood clot VCCE would allow direct comparison to these proposed models in the future

### 4.3. Critical values

In contrast to the constitutive model-based results discussed above, the critical pressure at onset of fracture (*P*_*crit*_) demonstrates observable rate dependence as shown in Figure 3A. As a ‘raw’ value obtained from an experiment, *P*_*crit*_ itself may be useful to characterize these materials regardless of the constitutive assumptions. The greater variability and increased median value of *P*_*crit*_ that we observe with faster strain rates points to open questions regarding the nature of this fracture transition and the rate dependence of fracture energy in whole blood clots.

### 4.4. Acceptability of using a miscible working fluid

Prior NICR and VCCE experimentation efforts in biological materials have largely avoided working fluids that could perfuse the sample being tested. Biological tissues, including clots, are known to be poroelastic under certain loading conditions (He et al., 2022; Oftadeh et al., 2018; Noailly et al., 2008). Most NICR studies avoid perfusion by using air, meanwhile, in controlled cavity expansion Nafo and Al-Mayah (2021) use a balloon to separate their sample from the cavity and Mijailovic et al. (2021) used silicone oil when testing in brain tissue. We chose PBS as a working fluid since it is biologically compatible, supporting the clinical relevance of our method, and resulted in more reliable operation of our test apparatus than oils or other fluids.

Using Darcy’s Law for pressure driven flow in porous media, we are able to show that fluid perfusion into the clot is insignificant on the timescales and pressure of our experiment. According to Darcy’s Law, the interstitial fluid flux (in m/s) is given as *q* = *kΔP/*(*μ*_*f*_ *L*) where *k* is the permeability, *ΔP* is the pressure drop that occurs over a distance *L*, and the dynamic viscosity is *μ*_*f*_. We simulate the fluid loss due to pressure driven flow by incrementally time stepping through a full cavity expansion process: first increasing the volume of the cavity, calculating the expected internal cavity pressure using the Ogden model, and then calculating fluid flux with Darcy’s Law. The cavity volume at the end of a time step is the initial volume plus injected volume less the lost volume calculated as flux through the surface area of the equivalent spherical cavity.

The permeability is taken as an upper bound seen in literature, and *L* is set as a single cavity radius (also an upper bound as the elastic response predicts a pressure gradient to occur over 10X the length). Wufsus et al. (2013) found the permeability *k* to range between 1.2×10^−1^ *μ*m^2^ and 1.5×10^−5^ *μ*m^2^ for platelet rich fibrin clots using perfusion testing with pressures between 30 and 2,000 kPa/m. He et al. (2022) used indentation and shear rheology to find a permeability of 3.5×10^−2^ *μ*m^2^ for human whole blood clots.

Results of this analysis, shown on Figure 6, indicate that no more than 2.5% of the injected fluid volume will be lost to interstitial flow under the experimental test conditions used in this work. Given that the expected fluid permeation is negligible and restricted to the vicinity of the cavity where pressure gradients are greatest, the permeation has little influence on the resisting pressure generated by the deformation field. Hence, we do not need to consider the coupling of poroelastic effects (as described by Biot (1956); Hu and Suo (2012) and others) with our measured material properties, and the incompressibility assumption that we employ in our theoretical model is valid.

**Figure 6:**
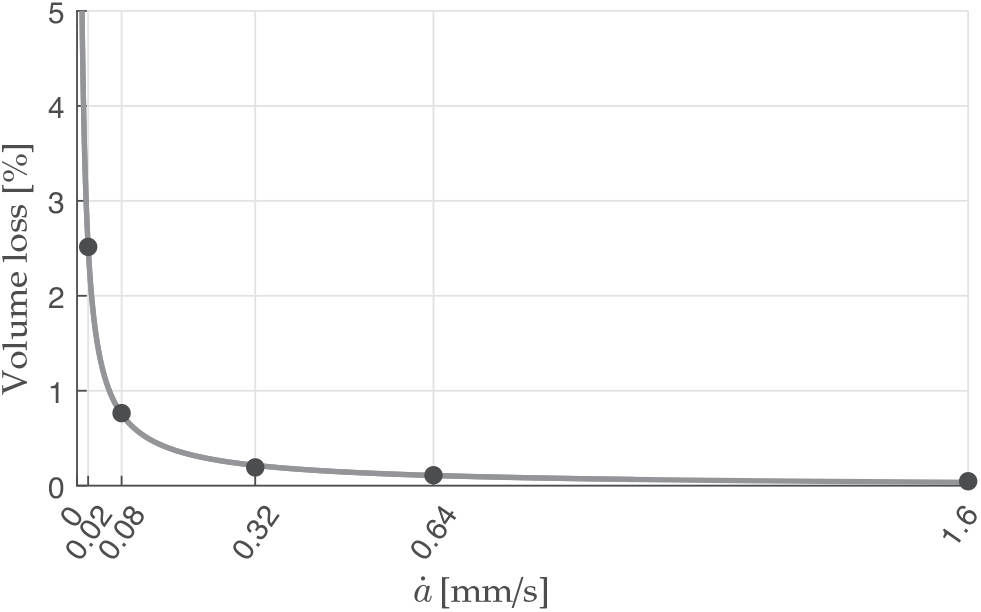
Fluid loss due to flow in the porous media is predicted to be less than 2.5% for the rates of expansion and cavity sizes considered in this test.

## 5. Conclusion

Thrombus dislodgement poses a significant risk to the population. Understanding the mechanical properties of whole blood clots is thus imperative for risk assessment as well as for the development of medical devices, therapies, and procedures that aim to safely remove clots or create clot mimics (Rausch et al., 2021a). Moreover, through development of *in silico* models, a deeper quantitative understanding can inform guidelines for safe transport loading after traumatic injury.

The application of VCCE to measure the elastic and rate dependent properties of whole blood clot is shown in this work. Performing radial expansion at rates from 0.02 to 1.60 mm/s and applying a 1-term Ogden model constitutive relationship we find that the shear modulus *μ* varies from 2.19 ± 0.15 to2.71 ± 0.78 kPa with a stiffening parameter of 5.69 ± 0.21 to 5.81 ± 0.31 across the tested rates. These results align with previous reported values for the nonlinear elastic properties of whole blood clot while providing more repeatable modulus measurements and larger samples sizes than the compression, tension, and simple shear methods described in literature.

We did not reproduce the strain rate dependence of stiffness that was previously reported for whole blood clots tested at slower rates, possibly demonstrating saturation of rate dependence at moderate strain rates. The expectation that the VCCE experimental procedure produces a consistent initial defect is validated by the repeatable initial defect size with *A* = 0.081 ± 0.011 mm, *N* = 36.

We have demonstrated that VCCE may be useful in other soft biological systems as well or in a clinical setting, given the relative ease of performing multiple, nondestructive and minimally invasive tests within the same sample and using saline as the working fluid. Our results demonstrate that VCCE can measure material properties with sufficient sensitivity to register elasticity changes as tissue ages, identifying a statistically significant difference in the elasticity of clots formed from blood stored for less than 3 days compared to blood stored for 3 or more days.

Furthermore, the precise volume control of the VCCE method provides the opportunity to extend investigations beyond the elastic regime discussed here, and into an investigation of fracture. Efforts to understand and model blood clot fracture are an active topic of research given the relevance to morbidity and mortality (Rausch et al., 2021b; Liu et al., 2021; Tutwiler et al., 2020). One avenue of future exploration is in understanding the observed dependence of *P*_*crit*_ and *a*_*crit*_ on strain rate. Numerous authors have begun investigating the transition between elastic cavitation or cavity expansion and fracture (Yang et al., 2019; Lefèvre et al., 2015; Kang et al., 2017; Hutchens et al., 2016; Lai et al., 2015; Song and Cai, 2022). Raayai-Ardakani et al. (2019b) have used energy minimization arguments in a VCCE setting to examine the critical values of transition to fracture and its propagation in PDMS, such approaches would have to be extended to capture the rate dependence observed in this work.

## Acknowledgements

We thank Prof. Benedetto Marelli and Dr. Eugene Lim of MIT for laboratory support. H.V. acknowledges the support of the Department of Defense (DOD) through the National Defense Science & Engineering Graduate (NDSEG) Fellowship Program. This material is based upon work supported by the National Science Foundation under Grant No. NSF-2127925, NSF-2046148, and NSF-2127925 to M.K.R. The authors further acknowledge the support of the Office of Naval Research under award number ONR-N00014-22-1-2073 to M.K.R. and ONR-N00014-20-1-2561 to T.C.. Funding agencies were not involved in study design; in the collection, analysis and interpretation of data; in the writing of the report; or in the decision to submit the article for publication.

## Footnotes

1 Sugerman et al. (2021b) have previously demonstrated coagulation times between 60 and 120 minutes do not have a significant effect on the stiffness of the resulting clot.

2 Earlier studies have shown that there is nearly no size sensitivity for *B/A >* 10, where *B* denotes the undeformed outer dimension of the body (Raayai-Ardakani et al., 2019a).

3 Note that this defect size corresponds to ∼20 − 25 red blood cells in diameter (Turgeon, 2005).

## References

Barney, C.W., Zheng, Y., Wu, S., Cai, S., Crosby, A.J., 2019. Residual strain effects in needle-induced cavitation. Soft Matter 15, 7390–7397.

Biot, M.A., 1956. Theory of deformation of a porous viscoelastic anisotropic solid. Journal of Applied physics 27, 459–467.

Budday, S., Sommer, G., Birkl, C., Langkammer, C., Haybaeck, J., Kohnert, J., Bauer, M., Paulsen, F., Steinmann, P., Kuhl, E., et al., 2017. Mechanical characterization of human brain tissue. Acta biomaterialia 48, 319–340.

Chockalingam, S., Roth, C., Henzel, T., Cohen, T., 2021. Probing local nonlinear viscoelastic properties in soft materials. Journal of the Mechanics and Physics of Solids 146, 104172.

Chueh, J., Wakhloo, A., Hendricks, G., Silva, C., Weaver, J., Gounis, M., 2011. Mechanical characterization of thromboemboli in acute ischemic stroke and laboratory embolus analogs. American Journal of Neuroradiology 32, 1237– 1244.

Cohen, T., Molinari, A., 2015. Dynamic cavitation and relaxation in incompressible nonlinear viscoelastic solids. International Journal of Solids and Structures 69, 544–552.

van Dam, E.A., Dams, S.D., Peters, G.W., Rutten, M., Schurink, G.W.H., Buth, J., van de Vosse, F.N., 2008. Non-linear viscoelastic behavior of abdominal aortic aneurysm thrombus. Biomechanics and modeling in mechanobiology 7, 127–137.

Gasser, T.C., Miller, C., Polzer, S., Roy, J., 2022. A quarter of a century biomechanical rupture risk assessment of abdominal aortic aneurysms. achievements, clinical relevance, and ongoing developments. International Journal for Numerical Methods in Biomedical Engineering, e3587.

Gersh, K.C., Nagaswami, C., Weisel, J.W., 2009. Fibrin network structure and clot mechanical properties are altered by incorporation of erythrocytes. Thrombosis and haemostasis 102, 1169–1175.

He, D., Kim, D.A., Ku, D.N., Hu, Y., 2022. Viscoporoelasticity of coagulation blood clots. Extreme Mechanics Letters 56, 101859.

Hinnen, J.W., Koning, O.H., Visser, M.J., Van Bockel, H.J., 2005. Effect of intraluminal thrombus on pressure transmission in the abdominal aortic aneurysm. Journal of vascular surgery 42, 1176–1182.

Hu, Y., Suo, Z., 2012. Viscoelasticity and poroelasticity in elastomeric gels. Acta Mechanica Solida Sinica 25, 441–458.

Huang, C.C., Chen, P.Y., Shih, C.C., 2013. Estimating the viscoelastic modulus of a thrombus using an ultrasonic shear-wave approach. Medical physics 40, 042901.

Hutchens, S.B., Fakhouri, S., Crosby, A.J., 2016. Elastic cavitation and fracture via injection. Soft Matter 12, 2557–2566.

Ji, Y., Dagro, A., Dorgant, G., Starr, D., Wilkerson, J., 2022. A comparison of conventional gel stiffness characterization techniques with cavitation rheology. Experimental Mechanics 62, 799–812.

Johnson, S., McCarthy, R., Gilvarry, M., McHugh, P.E., McGarry, J.P., 2021. Investigating the mechanical behavior of clot analogues through experimental and computational analysis. Annals of Biomedical Engineering 49, 420– 431.

Kang, J., Wang, C., Cai, S., 2017. Cavitation to fracture transition in a soft solid. Soft Matter 13, 6372–6376.

van Kempen, T.H., Donders, W.P., van de Vosse, F.N., Peters, G.W., 2016. A constitutive model for developing blood clots with various compositions and their nonlinear viscoelastic behavior. Biomechanics and modeling in mechanobiology 15, 279–291.

Krasokha, N., Theisen, W., Reese, S., Mordasini, P., Brekenfeld, C., Gralla, J., Slotboom, J., Schrott, G., Monstadt, H., 2010. Mechanical properties of blood clots–a new test method. Materialwissenschaft und Werkstofftechnik 41, 1019–1024.

Kundu, S., Crosby, A.J., 2009. Cavitation and fracture behavior of polyacrylamide hydrogels. Soft Matter 5, 3963–3968.

Lai, C.Y., Zheng, Z., Dressaire, E., Wexler, J.S., Stone, H.A., 2015. Experimental study on penny-shaped fluid-driven cracks in an elastic matrix. Proceedings of the Royal Society A: Mathematical, Physical and Engineering Sciences 471, 20150255.

Lefèvre, V., Ravi-Chandar, K., Lopez-Pamies, O., 2015. Cavitation in rubber: an elastic instability or a fracture phenomenon? International Journal of Fracture 192, 1–23.

Liang, X., Chernysh, I., Purohit, P.K., Weisel, J.W., 2017. Phase transitions during compression and decompression of clots from platelet-poor plasma, platelet-rich plasma and whole blood. Acta Biomaterialia 60, 275–290.

Liebeskind, D.S., Sanossian, N., Yong, W.H., Starkman, S., Tsang, M.P., Moya, A.L., Zheng, D.D., Abolian, A.M., Kim, D., Ali, L.K., et al., 2011. Ct and mri early vessel signs reflect clot composition in acute stroke. Stroke 42, 1237–1243.

Litvinov, R.I., Weisel, J.W., 2016. What is the biological and clinical relevance of fibrin? Seminars in thrombosis and hemostasis 42, 333–343.

Litvinov, R.I., Weisel, J.W., 2017. Role of red blood cells in haemostasis and thrombosis. ISBT science series 12, 176–183.

Liu, S., Bao, G., Ma, Z., Kastrup, C.J., Li, J., 2021. Fracture mechanics of blood clots: Measurements of toughness and critical length scales. Extreme Mechanics Letters 48, 101444.

Lohr, M.J., Sugerman, G.P., Kakaletsis, S., Lejeune, E., Rausch, M.K., 2022. An introduction to the ogden model in biomechanics: benefits, implementation tools and limitations. Philosophical Transactions of the Royal Society A 380, 20210365.

Lu, Y.C., Kemper, A.R., Untaroiu, C.D., 2014. Effect of storage on tensile material properties of bovine liver. Journal of the mechanical behavior of biomedical materials 29, 339–349.

Malone, F., McCarthy, E., Delassus, P., Fahy, P., Kennedy, J., Fagan, A., Morris, L., 2018. The mechanical characterisation of bovine embolus analogues under various loading conditions. Cardiovascular Engineering and Technology 9, 489–502.

Marder, V.J., Chute, D.J., Starkman, S., Abolian, A.M., Kidwell, C., Liebeskind, D., Ovbiagele, B., Vinuela, F., Duckwiler, G., Jahan, R., et al., 2006. Analysis of thrombi retrieved from cerebral arteries of patients with acute ischemic stroke. Stroke 37, 2086–2093.

Mijailovic, A.S., Galarza, S., Raayai-Ardakani, S., Birch, N.P., Schiffman, J.D., Crosby, A.J., Cohen, T., Peyton, S.R., Van Vliet, K.J., 2021. Localized characterization of brain tissue mechanical properties by needle induced cavitation rheology and volume controlled cavity expansion. Journal of the Mechanical Behavior of Biomedical Materials 114, 104168.

Nafo, W., Al-Mayah, A., 2021. Measuring the hyperelastic response of porcine liver tissues in-vitro using controlled cavitation rheology. Experimental Mechanics 61, 445–458.

Nickerson, C.S., Park, J., Kornfield, J.A., Karageozian, H., 2008. Rheological properties of the vitreous and the role of hyaluronic acid. Journal of Biomechanics 41, 1840–1846.

Noailly, J., Van Oosterwyck, H., Wilson, W., Quinn, T.M., Ito, K., 2008. A poroviscoelastic description of fibrin gels. Journal of biomechanics 41, 3265–3269.

Oftadeh, R., Connizzo, B.K., Nia, H.T., Ortiz, C., Grodzinsky, A.J., 2018. Biological connective tissues exhibit viscoelastic and poroelastic behavior at different frequency regimes: Application to tendon and skin biophysics. Acta biomaterialia 70, 249–259.

Ogden, R.W., 1972. Large deformation isotropic elasticity–on the correlation of theory and experiment for incompressible rubberlike solids. Proceedings of the Royal Society of London. A. Mathematical and Physical Sciences 326, 565–584.

Raayai-Ardakani, S., Chen, Z., Earl, D.R., Cohen, T., 2019a. Volume-controlled cavity expansion for probing of local elastic properties in soft materials. Soft matter 15, 381–392.

Raayai-Ardakani, S., Cohen, T., 2019. Capturing strain stiffening using volume controlled cavity expansion. Extreme Mechanics Letters 31, 100536.

Raayai-Ardakani, S., Earl, D.R., Cohen, T., 2019b. The intimate relationship between cavitation and fracture. Soft matter 15, 4999–5005.

Rausch, M.K., Parekh, S.H., Dortdivanlioglu, B., Rosales, A.M., 2021a. Synthetic hydrogels as blood clot mimicking wound healing materials. Progress in Biomedical Engineering 3, 042006.

Rausch, M.K., Sugerman, G.P., Kakaletsis, S., Dortdivanlioglu, B., 2021b. Hyper-viscoelastic damage modeling of whole blood clot under large deformation. Biomechanics and Modeling in Mechanobiology 20, 1645–1657.

Riha, P., Wang, X., Liao, R., Stoltz, J., 1999. Elasticity and fracture strain of whole blood clots. Clinical hemorheology and microcirculation 21, 45–49.

Song, Z., Cai, S., 2022. Needle-induced-fracking in soft solids with crack blunting. Extreme Mechanics Letters 52, 101673.

Sugerman, G.P., Chokshi, A., Rausch, M.K., 2021a. Preparation and mounting of whole blood clot samples for mechanical testing. Current Protocols 1, e197.

Sugerman, G.P., Kakaletsis, S., Thakkar, P., Chokshi, A., Parekh, S.H., Rausch, M.K., 2021b. A whole blood thrombus mimic: Constitutive behavior under simple shear. Journal of the Mechanical Behavior of Biomedical Materials 115, 104216.

Sugerman, G.P., Parekh, S.H., Rausch, M.K., 2020. Nonlinear, dissipative phenomena in whole blood clot mechanics. Soft Matter 16, 9908–9916.

Turgeon, M.L., 2005. Clinical hematology: theory and procedures. Lippincott Williams & Wilkins.

Tutwiler, V., Mukhitov, A.R., Peshkova, A.D., Le Minh, G., Khismatullin, R., Vicksman, J., Nagaswami, C., Litvinov, R.I., Weisel, J.W., 2018. Shape changes of erythrocytes during blood clot contraction and the structure of polyhedrocytes. Scientific reports 8, 1–14.

Tutwiler, V., Singh, J., Litvinov, R.I., Bassani, J.L., Purohit, P.K., Weisel, J.W., 2020. Rupture of blood clots: Mechanics and pathophysiology. Science advances 6, eabc0496.

Tynngård, N., Lindahl, T., Ramstrom, S., Berlin, G., 2006. Effects of different blood components on clot retraction analysed by measuring elasticity with a free oscillating rheometer. Platelets 17, 545–554.

Umale, S., Deck, C., Bourdet, N., Dhumane, P., Soler, L., Marescaux, J., Willinger, R., 2013. Experimental mechanical characterization of abdominal organs: liver, kidney & spleen. Journal of the mechanical behavior of biomedical materials 17, 22–33.

Weafer, F.M., Duffy, S., Machado, I., Gunning, G., Mordasini, P., Roche, E., McHugh, P.E., Gilvarry, M., 2019. Characterization of strut indentation during mechanical thrombectomy in acute ischemic stroke clot analogs. Journal of neurointerventional surgery 11, 891–897.

World Health Organization, 2020. The top 10 causes of death. URL: https://www.who.int/news-room/fact-sheets/detail/the-top-10-causes-of-death. accessed: 6 Oct 2022.

Wufsus, A.R., Macera, N., Neeves, K., 2013. The hydraulic permeability of blood clots as a function of fibrin and platelet density. Biophysical journal 104, 1812–1823.

Yang, S., Bahk, D., Kim, J., Kataruka, A., Dunn, A.C., Hutchens, S.B., 2019. Hydraulic fracture geometry in ultrasoft polymer networks. International Journal of Fracture 219, 89–99.

Zimberlin, J.A., McManus, J.J., Crosby, A.J., 2010. Cavitation rheology of the vitreous: mechanical properties of biological tissue. Soft Matter 6, 3632– 3635.

Zimberlin, J.A., Sanabria-DeLong, N., Tew, G.N., Crosby, A.J., 2007. Cavitation rheology for soft materials. Soft Matter 3, 763–767.

