## Supplemental Information 1 for "Elasticity of Whole Blood Clots Measured via Volume Controlled Cavity Expansion"

### Electronic Supplemental Information for: Elasticity of Whole Blood Clots Measured via Volume Controlled Cavity Expansion

Hannah Varner, Gabriella P. Sugerman, Manuel K. Rausch, Tal Cohen

---

---

#### ESI 1. Example fitting curves

The Ogden model captures the strain stiffening elastic response of blood clots across test rates and days. Fitting is performed as described in Section 2.4 of the manuscript.

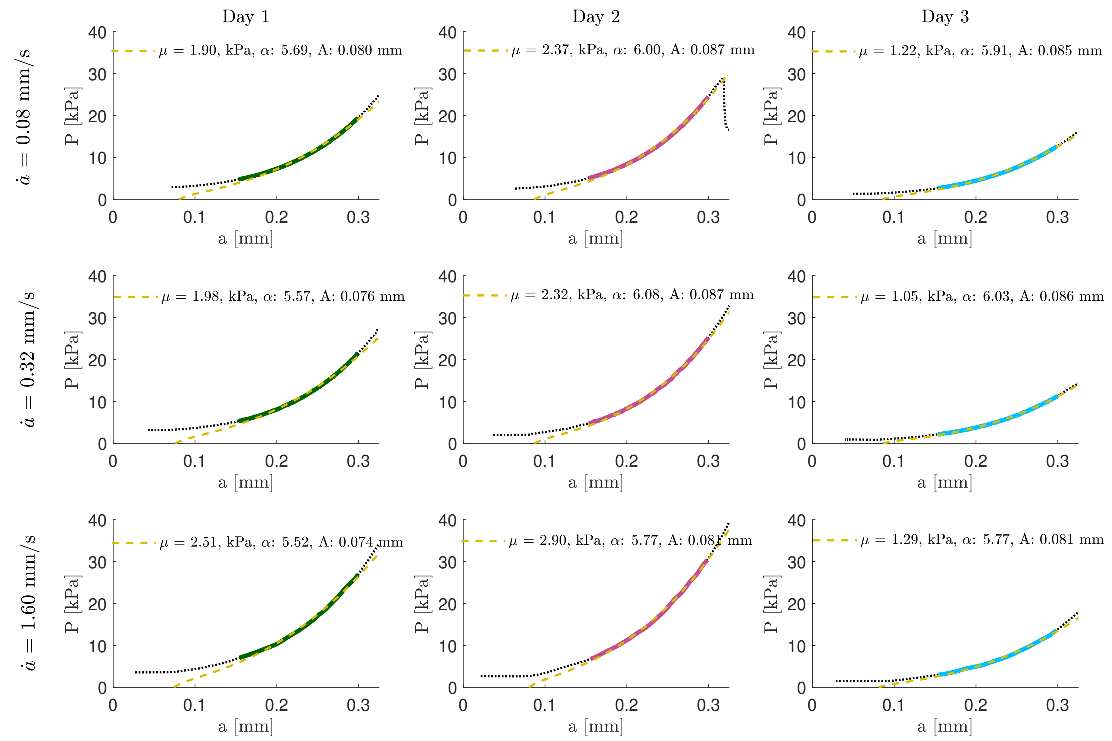


Figure A1: Representative fitted curves for individual tests. Black dotted line is the full set of experimental data, colored region is the truncated data between $a_{min}$ and $a_{max}$ used for fitting, and the yellow dashed line is the fit resulting from our NLLS optimization.
